# Insights on the effect of extracellular acidification on an HCN channel: a molecular dynamics study

**DOI:** 10.1101/2025.09.05.674486

**Authors:** Somayeh Asgharpour, Michalis Lazaratos, Marc Spehr, Paolo Carloni

## Abstract

Extracellular acidification may affect the structure and function of ion channels by neutralizing acidic residues exposed to the extracellular space. However, the relationship between these changes in the protonation state and alterations in the channels’ structure and dynamics remains unclear. Here, we used atomistic simulations and graph-based algorithms to study mice hyperpolarization-activated cyclic nucleotide-gated type 2 channels, whose gating is facilitated by extracellular acidification. Our simulations revealed that E212, a residue facing the extracellular space, may be involved in complex hydrogen bond networks with the S4 helix, which plays a role in the gating mechanism. This network is partially disrupted at acidic pH levels, affecting the molecular interactions of the S4 helix. This, in turn, may alter the domain’s response to membrane voltage changes and, consequently, gating. We conclude that the H-bond network from an extracellular residue to the main gating domain may be an important factor in the observed channel activities at different pH levels.

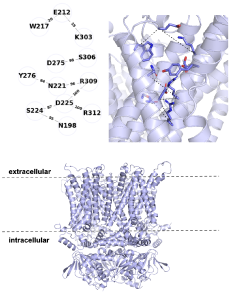

## 1 Introduction

Extracellular acidification may affect the voltage-dependent gating or conduction in ion channels, including selective potassium [1–8], sodium [9] [10], and chloride channels [11], along with non-selective ones such as hyperpolarization and cyclic nucleotide activated (HCN) [12] and acid-sensing ion channels [13].

The properties of the ion channel can change as a result of altered protonation states, which in turn can affect intermolecular hydrogen bonds that originate at the extracellular site and extend into the protein interior. Thus, changes in the water-wired H-bond networks and long-distance H-bond pathways (which can interconnect remote protein domains) might influence the gating process. Here, we address this issue by performing molecular dynamics (MD) simulations and graph-based H-bond analysis [14] [15] on extracellular acidification in a specific channel, mouse hyperpolarization-activated cyclic nucleotide-gated ion channel type 2 (HCN2) [16]. This acidification leads to a positive shift in the voltage dependence of activation [12], which facilitates the gating mechanism [17].

The HCN2 channel consists of four identical subunits (Fig. 1a). Each of them includes six transmembrane *α* helices (S1-S6), the *pore domain* and the *HCN domain*, which is a bundle of three helices specific to these channels^1^. The transmembrane domain in turn interacts non-covalently with the cyclic nucleotide-binding domain (CNBD) [18] (Fig. 1). Activation of the channel and ion conduction relies on the collective motion of helices S1-S4 (*voltage sensing domain (VSD)*) and the pore. For the pore to open, this region must undergo a significant conformational change (Fig. 1e) [19], associated with a rearrangement of salt bridges and H-bond interactions between the nine basic residues of S4 (Fig. 1e) and the polar and negatively charged residues of the helices S2 and S3 [20] [21] (Fig. 1e). The key question addressed here is whether this network of interactions changes with extracellular acidification in the inactive state and whether such changes may affect the gating process.

**Figure 1:**
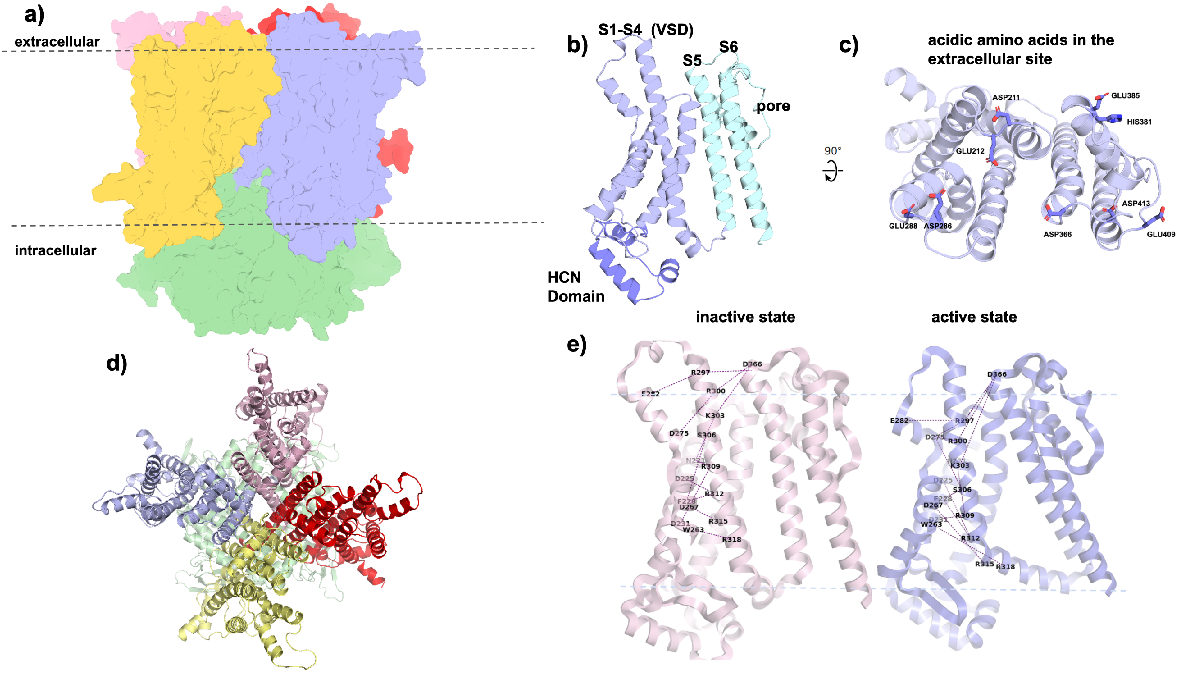
The mouse HCN2 structure was predicted based on the highly similar human HCN1 X-ray structure (PDB code: 5U6O [22]), which shares 91% sequence identity (Fig. S1). **(a)** Structural model of the mouse HCN2 channel. Each subunit is shown in a different color; the cyclic nucleotide-binding (CNB) domain is highlighted in green. **(b)** The subunits contain -helices S1–S6, the pore, and the HCN domain, which forms non-covalent interactions with the CNB domains. **(c)** The titratable extracellular amino acids in one subunit. **(d)** Orientation of the channel showing the pore and extracellular site. **(e)** Comparison of the active-state structure (PDB code: 6UQF [19]) with the inactive state, highlighting differences in the hydrogen-bonding pattern.

## 2 Methods

### Structural models

A model of the mouse HCN2 channel including all domains (VSD, CNBD, HCND, and the pore domain) was generated by homology modeling using the structure of human HCN1 (see SI for details). The *pK*_*a*_ values of the extracellular residues were calculated using PROPKA (Tab. S1) [23] [24] and the protonation states were adjusted accordingly.

### MD simulations

We used the CHARMM36 force field for the protein, lipids, and ions [25] and the TIP3P water model [26]. Long-range interactions were evaluated using particle-mesh Ewald (PME) summation with a cutoff of 12 Å for the real-space component. The Lennard-Jones interactions were truncated at 12 Å with an atom-based force-switching function, effective at 10 Å. The integration time step was set to 2 fs. All bonds were constrained by the LINCS algorithm [27]. After an equilibration phase, two 1.2 *µ*-s long NPT simulations were performed using the Nosé-Hoover thermostat [28] with an effective relaxation time of 1 ps, a semi-isotropic Parrinello-Rahman barostat [29] and a pressure coupling effective every 5 ps with a compressibility of 4.5 × 10^*−*5^. The last 800 ns of each simulation were used for analysis. All MD simulations were performed using the GROMACS-2020 package [30].

### Graph-based analyses

Graphs of water-mediated and/or direct H-bond networks were generated using the Bridge code [14] and the Bridge2 graphical user interface [31] (see SI for details). Here, we computed these graphs using 8,000 equally spaced coordinate snapshots from the last 800 ns of each simulation, with time intervals of 100 ps. H-bonds with high occupancies > 50% contribute to the analysis. Additional details and validation procedures are provided in the SI.

## 3 Results and Discussion

Each of the four subunits, at the extracellular side, contains one Histidine, two Aspartate and six Glutamate residues (H381, E385, D366, E409, D413, D211, E212, D286, and E288) as titratable residues. At physiological pH, the histidine residue is assumed to be neutral and protonated on its *δ* Nitrogen, while the acidic residues are assumed to be negatively charged. At pH 3.0 (in the extracellular medium) these residues are all protonated, while the H381 is positively charged (see Tab. S1).

Two 1.2 *µ*s-long MD simulations of the channel, including membrane and solvents and ions at physiological pH and upon extracellular acidification at pH 3.0 were performed and well maintained during the dynamics (see SI for details). Both simulations were converged after 0.4 *µ*s, as from RMSD analysis (Fig. S2).

Next, we present the results for one subunit.

At pH 7.0 a rather persistent H-bond network D275-R309-D225-R312 (occupancy = 100%) is present inside the VSD which maintains the channel in its closed conformation. E212 connects to this group of residues, by water mediated interactions either through Y276, or D275, or W217 (Fig. 2a, 2b). Specifically, we find the following pathways at pH 7.0: **path1**: E212-Y276-D275-R309-D225-R312 (Fig. 3a), with an occupancy threshold of 80% (Fig. 2a). **path2**: E212-D275-R312 (Fig. 3b) with an occupancy threshold of 53% (Fig. 3a), and **path3**: E212-W217-R309-D225-R312 (Fig. 3c) with an occupancy threshold of 55% (Fig. 3a). These networks involve as many as 3 waters involved in each water wired bond. E212 forms also a salt bridge with K303 (Fig. 2a,2d).

**Figure 2:**
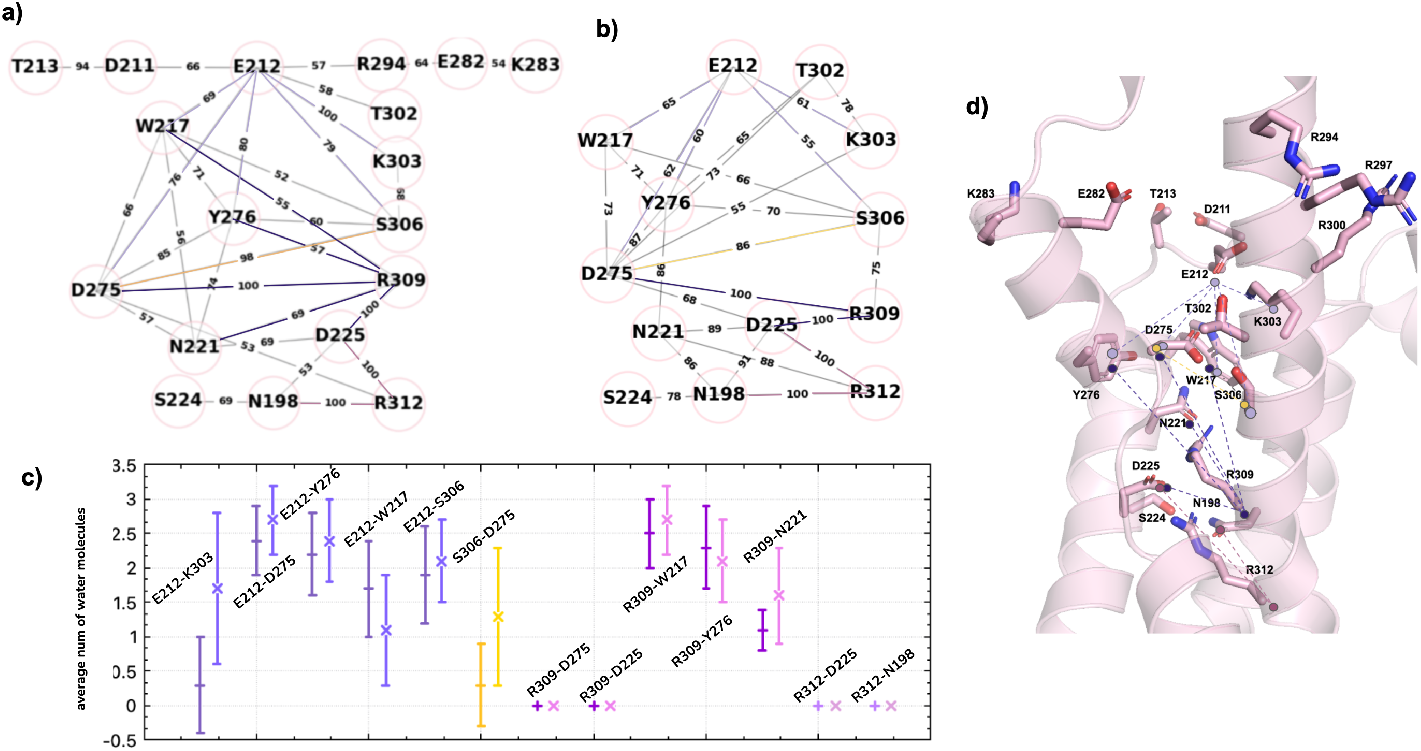
**(a)** A highly persistent H-bond network connects E212 to R312 in the VSD at pH 7.0. **(b)**. At pH 3.0, the network is partly disrupted. Key interactions are shown with colored lines. **(c)** The average number of water molecules among residues. Coloring scheme as in (a) and (b). **(d)** Structure of the portion of the protein involved in the network. Key aminoacids are shown here with the same color as in the network

**Figure 3:**
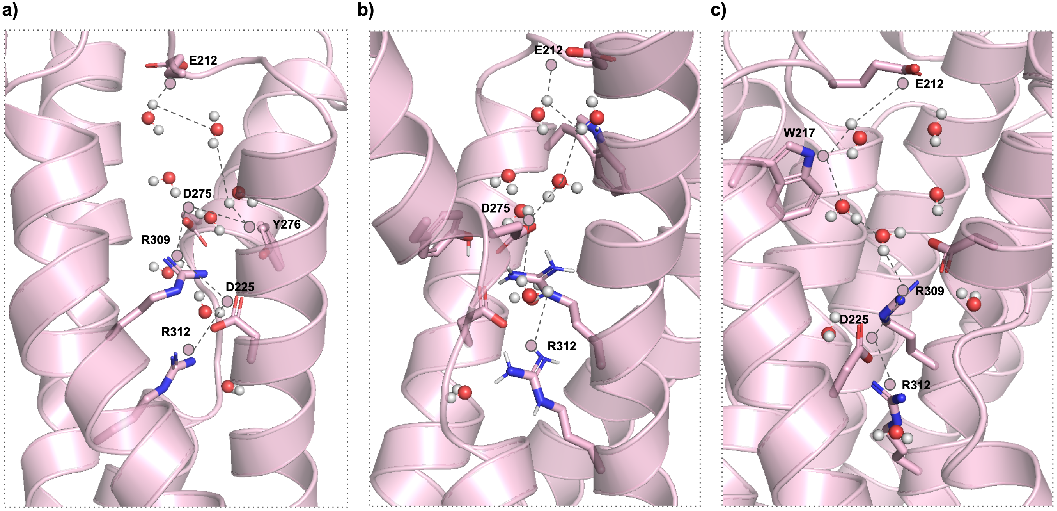
Three water mediated pathways that connect E212 to R312

At pH 3.0 (Fig. 2b), the occupancy of the salt bridge between E212 and K303 is reduced from 100% to 60%. This may be expected because protonated E212 does not form a salt bridge with K303 but only H-bond interactions (Fig. 2d). Consequently, protonated E212 is much more mobile than at pH 7.0. As a result, **Path1** is present and E212 interacts with Y276 with a lower occupancy 60%; the other two pathways are absent due to the disruption of the water-mediated H-bond interactions between E212 with either D275 on **path2** or W217 on **path3**. The other extracellular residues’ H-bond networks are not affected b y a cidification (d ata not shown).

The results of the other subunits are rather similar (see SI) except that in one subunit an adjacent residue (T213) replaces E212 in forming an H-bond network with the VSD. Thus, the effect we s ee h ere m ay for one subunit may be present also in others.

Taken together, these simulations may provide a first insight on the observed effect of acidification on gating at the molecular scale [12]. The water-mediated H-bond network E212-R312@S4 weakens dramatically at low pH, likely affecting the interactions of S4 and its enviroment [19]: The helix becomes more “loose” at pH 3.0. The effect is observed also in other subunits. Thus, S4 motion toward the cytoplasm - one of the hallmarks of the gating process [19]-might be facilitated, as experimentally observed [12]. Mutation of E212 into a non-ionizable group might therefore impact the pH dependence of the gating. Free energy calculations of the gating process at different pH values are required to test this hypothesis.

## Supporting information

https://it.overleaf.com/8895148658dknjrcjsvmpv#d06c88

## Author information

SA performed all simulations, calculations, and analyses. PC supervised the study; MS and PC acquired the fundings; SA drafted the original manuscript; MS, PC, revised the original draft; ML contributed to early H-bond graph analyses and discussions and guidance. All authors read and corrected the manuscript.

## Data availability statement

Data supporting the findings of this study are available from the first author.

## Acknowledgments

The authors thank (i) the GRK 2416 project (MultiSenses-MultiScales: Novel approaches to decipher neural processing in multisensory integration) for funding this project (grant reference number 368482240/GRK2416). (ii) the Juelich Super Computer Center for computer time. (iii) Dr. America Luz Chi Uluac for fruitful discussions, Dr Ana-Nicoleta Bondar for support with the Bridge Code, and Dr. Emiliano Ippoliti for the use of the HPC facilities in Jülich.

## Footnotes

1 This Domain is not present in other voltage-gated channels [18]

