## Supplementary material for "Insights on the effect of extracellular acidification on an HCN channel: a molecular dynamics study": https://it.overleaf.com/8895148658dknjrcjsvmpv#d06c88

### Supporting Information
### Insights on the effect of extracellular acidification on anHCN channel

Somayeh Asgharpour[1,2], Michalis Lazaratos[3], Marc Spehr[1,4], and Paolo Carloni[1,2]

[1]Research Training Group 2416 MultiSenses – MultiScales, RWTH Aachen University, Aachen, Germany
[2]IAS-5/INM-9, Computational Biomedicine, Forschungszentrum Jülich, Wilhelm-Johnen-Strasse, 52428 Jülich, Germany
[3]Department of Physics, Theoretical Molecular Biophysics Group,Freie Universität Berlin, Berlin, Germany
[4]Department of chemosensation, Institute for biology II, RWTH Aachen University, Aachen, Germany

September 5, 2025

### 1 Computational details

#### 1.1 Protonation states

We used the **Propka 3.1** module [1] [2] in Maestro Schrödinger software [3] to obtain estimated values $pK_a$ of ionizable residues at the extracellular site in the resting state of the channel (Table S1).

| residue | $pK_a$ |
|---|---|
| ASP 211 | 3.10 |
| GLU 212 | 5.29 |
| ASP 286 | 3.93 |
| GLU 288 | 4.57 |
| ASP 366 | 4.01 |
| ASP 370 | 3.99 |
| GLU 385 | 3.01 |
| GLU 409 | 4.91 |
| ASP 413 | 5.28 |

Table S1: Estimated $pK_a$ values of ionizable residues located at the extracellular site of the channel.

#### 1.2 Homology modeling of the mouse HCN2 channel

The human HCN1 channel, for which a cryo-EM structure in the close conformation has been solved (PDB code 5u6o) and shares 91% sequence identity with the mouse HCN2 channel (UniProtKB – O88703 (HCN2-MOUSE); https://www.uniprot.org/uniprot/O88703)(Fig. S1) was used as a template to derive the mouse HCN2 structure in the closed conformation, using the Swiss model [4].

```
                              HCN-Domain
Mice   136  LQRQFGALLQPGVNKFSLRMFGSQKAVEREQERVKSAGAWIIHPYSDFRFYWDFTMLLFM   195
Human   94  MQRQFTSMLQPGVNKFSLRMFGSQKAVEKEQERVKTAGFWIIHPYSDFRFYWDLIMLIMM   153
                  S1                       S2
Mice   196  VGNLIIIPVGITFFKDETTAPWIVFNVVSDTFFLMDLVLNFRTGIVIEDNTEIILDPEKI   255
Human  154  VGNLVIIPVGITFFTEQTTTPWIIFNVASDTVFLLDLIMNFRTGTVNEDSSEIILDPKVI   213
                              S3                              S4
Mice   256  KKKYLRTWFVVDFVSSIPVDYIFLIVEKGIDSEVYKTARALRIVRFTKILSLLRLLRLSR   315
Human  214  KMNYLKSWFVVDFISSIPVDYIFLIVEKGMDSEVYKTARALRIVRFTKILSLLRLLRLSR   273
                                           S5
Mice   316  LIRYIHQWEEIFHMTYDLASAVMRICNLISMMLLLCHWDGCLQFLVPMLQDFPSDCWVSI   372
Human  274  LIRYIHQWEEIFHMTYDLASAVVRIFNLIGMMLLLCHWDGCLQFLVPLLQDFPPDCWVSL   333
                             H5                               S6
Mice   373  NNMVNHSWSELYSFALFKAMSHMLCIGYGRQAPESMTDIWLTMLSMIVGATCYAMFIGHA   429
Human  334  NEMVNDSWGKQYSYALFKAMSHMLCIGYGAQAPVSMSDLWITMLSMIVGATCYAMFVGHA   393

Mice   430  TALIQSL   437
Human  394  TALIQSL   400
```

Figure S1: Sequence alignment between human HCN1 (5u6o) and Mice HCN2

The amino acid sequence of the S4 helix shows a sequence identity of 100% with human HCN1. For S1, the sequence identity is 82%, for S2, 70%; and for S3 79%. The helices show some variations as in S1, changes from hydrophobic to polar (I $\rightarrow$ T) and aliphatic to aromatic (L $\rightarrow$ F). In S2, we observe changes in aromaticity (V $\rightarrow$ F). In S3 on the other hand, we observe an increase in the charge and hydrophobic to basic shifts (V $\rightarrow$ K, M $\rightarrow$ K). The loop connecting the S1 and S2 helices that contains D211 and E212 is altered (TEQ $\rightarrow$ KDE) so E212 is an additional charged amino acid in the mice isoform not present in human HCN1. Furthermore, the loop that connects the pore and S5 includes a histidine, replacing an aspartate found in the human isoform. Site-directed mutagenesis studies of the pore domain demonstrated alterations that leads to changes in ion permeability [5]. Many mutations are reported for the S4 helix that affect the channel function where they alter positively charged amino acids (R $\rightarrow$ Q, K $\rightarrow$ Q) which shifts the voltage dependence to more negative values [6]. For the loop that we focus on in this study, T $\rightarrow$ R or T $\rightarrow$ P mutations have been reported in Human HCN1 with uncertain significance [7].

### 2 Methods

#### 2.1 MD simulations

The systems were embedded in a hydrated bilayer of palmitoyl-oleoyl-sn-glycero-phosphocholine (POPC) lipids using the CHARMM-GUI server [8]. They contained ≈ 70,000 a water molecules. The system at pH 7.0 contained 190 $K^+$ and 198 $Cl^-$ ions, while the one at pH 3.0 included 190 $K^+$ and 238 $Cl^-$ ions. In this way, both systems were electrically neutral in ≈ 150mM KCl. They consisted of ≈ 300,000 atoms, embedded in a simulation box of edges ≈ 130 Å, 130 Å, 170 Å.

The systems were minimized after 2,000 steps of steepest descent. Each system was then equilibrated for 1 ns in the NVT ensemble (T = 300K). This was followed by a 1 ns-long equilibration in the NPT ensemble ( P = 1 atm ). We used Berendsen's thermostat and barostat [9]. The production run was then performed for 1.2 $\mu$s for each system keeping the temperature at 300 K by Nosé-Hoover thermostat and the pressure at 1 atm by parinello-Rahman barostat.

#### 2.2 Analysis of the MD simulations

The Root-Mean-Square-Deviation (RMSD) is shown for the four VSD helices with and without the connecting loops. The value ranges between 0.0-5.0 Å. It is stabilized after almost 400 ns (Fig. S2).

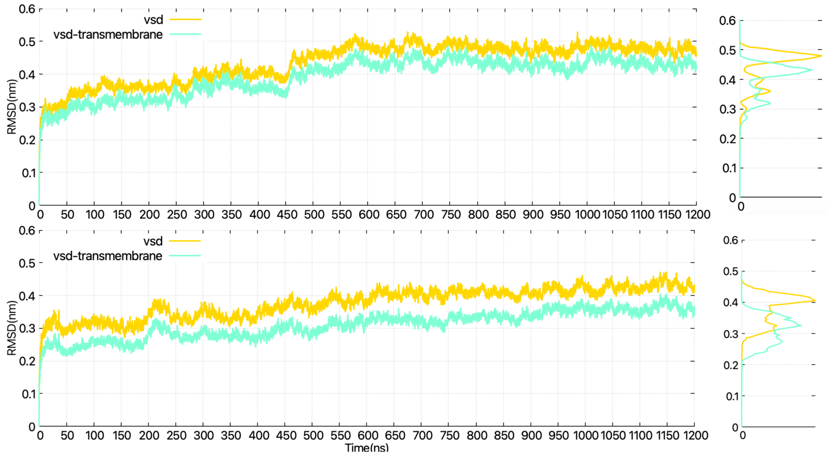


Figure S2: The RMSD profile of the VSD in both simulations pH 7.0 (up) and pH 3.0 (down) , smoothened histograms of the rmsd profiles for the entire trajectory

The Root-Mean-Square-Fluctuations (RMSF) of the VSD and the connecting loops are reported in Fig. S3. The highest fluctuations belong to the loops connecting S2 and S3 to each other. In subunit C, the RMSF is higher at pH 7.0 than at pH 3.0 (see Fig. S3), while in all other subunits, they are either in the same range or lower at pH 7.0 so the fluctuation is reduced upon protonation. The loop connecting the S2 and S3 helix fluctuates more at lower pH in subunit A, less in subunit C, and is the same range in subunits B and D. E212 shows a higher RMSD in subunit C once it is deprotonated while in other subunits it was either in the same range or lower.

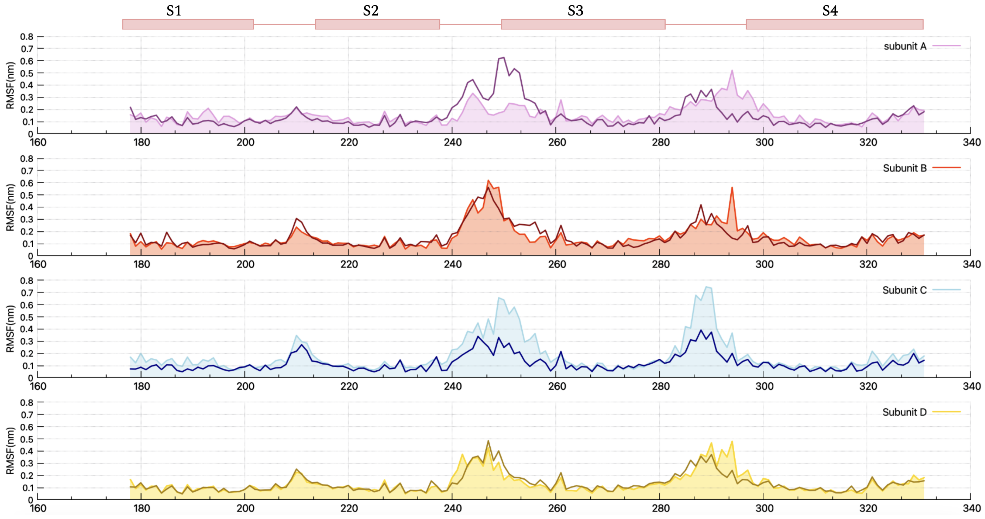


Figure S3: RMSF of the residues of the VSD and its connecting loops in both simulations; the light colors refer to the simulation at pH 7.0 and darker colors refer to that at pH 3.0

### 3 H-Bond network analysis

An H-bond is assumed to be formed when the distance between the H-bond donor and acceptor is within 3.5 Å, and the H-bond angle is within 60 degrees. An occupancy of H-bond of 100% means that the H-bond is formed throughout the entire trajectory segment used for data analysis.

#### 3.1 The S4 helix

The main interactions mediating the key motion in the S4 helix [10] are shown in Fig. S4a. Fig. S4b compares their occupancy values in 4 subunits and both simulations. Lines indicate occupancies at pH 7.0, and dashed lines indicate occupancies at pH 3.0. Fig. S4c shows that keeping the network at an occupancy cutoff of more than 75% maintains most of these interactions and possible pathways from E212 to R312 and accurately reflects our aim.

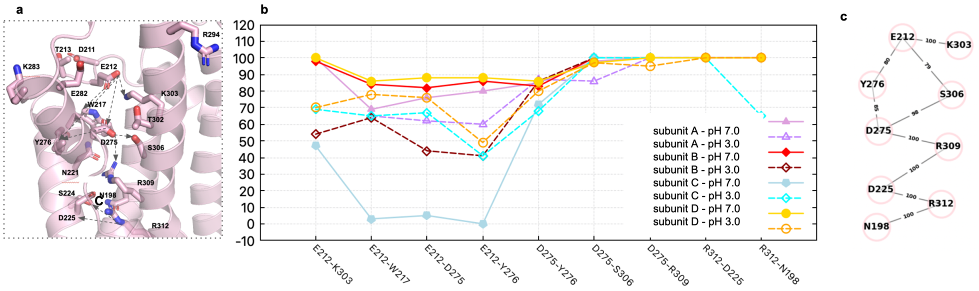

Figure S4: (a) H-bond pairs that in the VSD. (b) Comparison of the occupancies of each H-bond in (a), for both simulations and for 4 subunits. (c) H-bond pairs with high occupancies shown as networks

As shown in Fig. S4b all subunits in both simulations had more than 90% occupancy when pairs involved R306, R309, and R312 with an exception for interactions with and N198 which have a 60% occupancy. However, pairs involving E212 showed a wider range of difference.

#### 3.2 Clustering Analysis

Fig. S5 shows the most common conformations of E212 at pH 7.0, as obtained by clustering analysis. In subunits B,D E212 is in contact with K303. Fig. S6 shows H-bond networks that connect E212 to R312 with an occupancy cut-off of 50%, similar to what was seen for subunit A (see main text). The networks feature fewer H-bonds and lower occupancies at pH 3.0. This supports the idea that protonation of E212 contributes to channel activation by disconnecting it from the VSD. In subunit C, T213 replaces E212 in forming an H-bond network with the VSD Fig. S6. In fact, E212 interacts with water molecules for more than 80% of the simulation time (Fig. S5) .

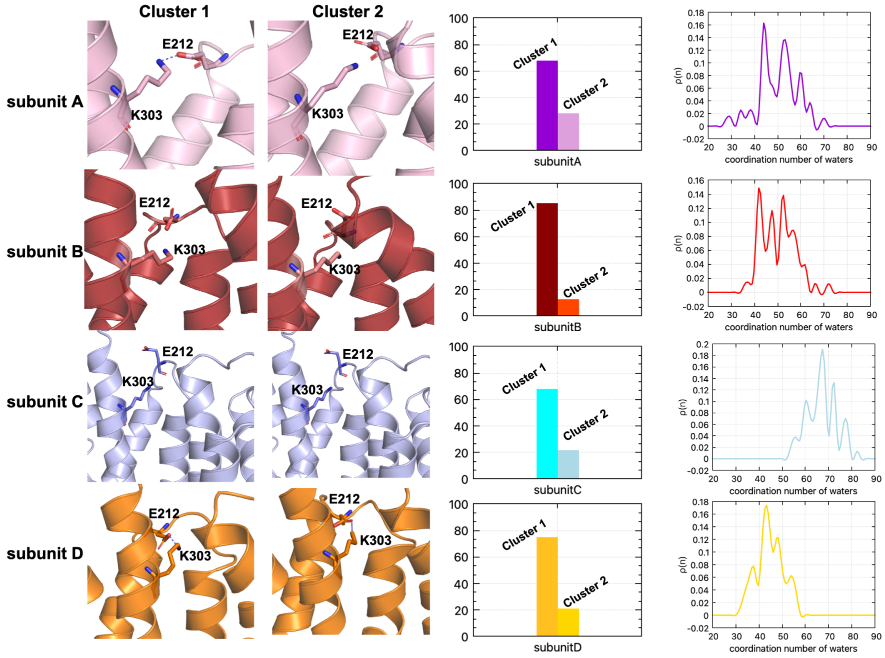


Figure S5: The largest clusters of E212 for each subunit in the MD trajectory at pH 7.0. Their populations are reported. The water coordination is also shown for each subunit where subunit C is more water exposed than subunits B, C, and D with a significant shift.

#### 3.3 H-bond networks

Here we report the H-bond netwroks of subunit B, C, and D obtained by similar conditions as subunit A reported in the main text.

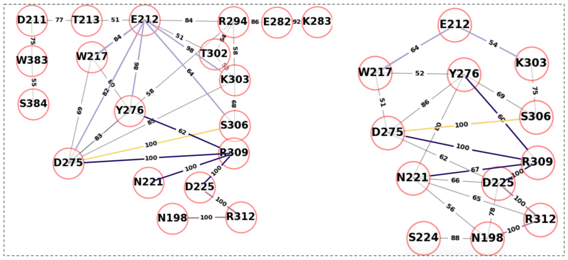


(a) networks compared for subunit B pH 7.0 (left), pH 3.0 (right)

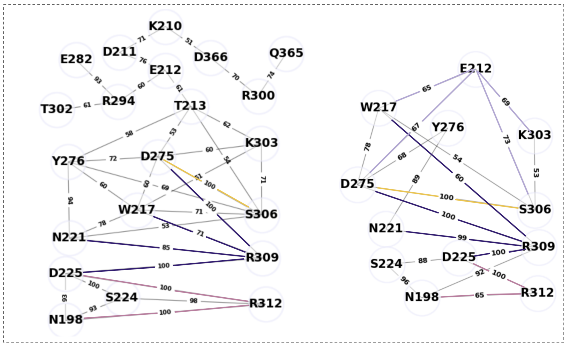


(b) networks compared for subunit C pH 7.0 (left), pH 3.0 (right)

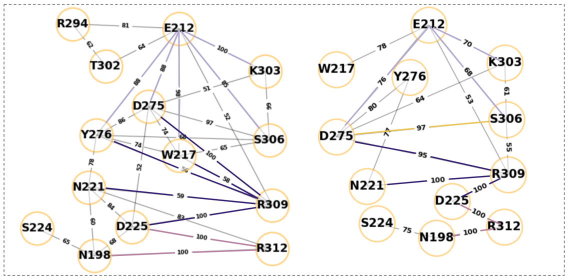


(c) networks compared for subunit D pH 7.0 (left), pH 3.0 (right)

Figure S6: H-bond network connecting residues at the extracellular side (E212, T213) of subunits B-D with the VSD

Tables S2, S3, and S4 report the number of waters involved in each H-bond network formed by E212 or T213. Similarly to subunit A, number of waters involved in E212-K303 are higher at pH 3.0 than at pH 7.0 with the exception of subunit C.

| Pairs | pH 7.0 | pH 3.0 |
|---|---|---|
| E212-K303 | 1.3 ∓ 0.9 | 1.6 ∓ 1.1 |
| E212-D275 | 2.2 ∓ 0.8 | 2.3 ∓ 0.5 |
| E212-Y276 | 2.3 ∓ 0.6 | 2.2 ∓ 0.5 |
| E212-W217 | 1.8 ∓ 0.7 | 0.4 ∓ 0.7 |
| E212-S306 | 2.4 ∓ 0.6 | 1.8 ∓ 0.8 |
| D275-S306 | 0.2 ∓ 0.4 | 0.1 ∓ 0.4 |
| R309-D275 | 0.4 ∓ 0.5 | 0.1 ∓ 0.4 |
| R309-D225 | 0.0 ∓ 0.0 | 0.0 ∓ 0.0 |
| R309-W217 | 2.4 ∓ 0.6 | 2.3 ∓ 0.5 |
| R309-Y276 | 1.5 ∓ 0.8 | 2.4 ∓ 0.6 |
| R309-N221 | 0.0 ∓ 0.1 | 1.0 ∓ 0.6 |
| R312-N198 | 0.0 ∓ 0.0 | 0.0 ∓ 0.0 |
| R312-D225 | 0.0 ∓ 0.0 | 0.0 ∓ 0.0 |

Table S2: Water analysis on subunit B

| Pairs | pH 7.0 | pH 3.0 |
|---|---|---|
| E212-K303 | 2.1 ∓ 1.0 | 1.3 ∓ 1.1 |
| E212-D275 | 2.9 ∓ 0.3 | 2.2 ∓ 0.5 |
| E212-Y276 | 2.7 ∓ 0.5 | 2.2 ∓ 0.7 |
| E212-W217 | 2.6 ∓ 0.6 | 1.1 ∓ 1.0 |
| E212-S306 | 2.9 ∓ 0.3 | 1.3 ∓ 0.7 |
| D275-S306 | 0.0 ∓ 0.2 | 0.0 ∓ 0.2 |
| R309-D275 | 0.0 ∓ 0.0 | 1.1 ∓ 0.3 |
| R309-D225 | 0.0 ∓ 0.0 | 0.0 ∓ 0.0 |
| R309-W217 | 2.0 ∓ 0.4 | 2.2 ∓ 0.4 |
| R309-Y276 | 2.5 ∓ 0.5 | 2.5 ∓ 0.5 |
| R309-N221 | 1.0 ∓ 0.1 | 0.0 ∓ 0.2 |
| R312-N198 | 0.0 ∓ 0.0 | 1.0 ∓ 0.2 |
| R312-D225 | 0.0 ∓ 0.0 | 0.0 ∓ 0.0 |

Table S3: Water analysis on subunit C

| Pairs | pH 7.0 | pH 3.0 |
|---|---|---|
| E212-K303 | 0.1 ∓ 0.3 | 1.2 ∓ 1.1 |
| E212-D275 | 2.3 ∓ 0.5 | 2.2 ∓ 0.5 |
| E212-Y276 | 2.1 ∓ 0.6 | 2.2 ∓ 0.7 |
| E212-W217 | 1.4 ∓ 0.7 | .5 ∓ 0.8 |
| E212-S306 | 1.9 ∓ 0.7 | 1.5 ∓ 0.8 |
| D275-S306 | 0.5 ∓ 0.8 | 0.2 ∓ 0.7 |
| R309-D275 | 0.0 ∓ 0.0 | 1.4 ∓ 0.7 |
| R309-D225 | 0.0 ∓ 0.0 | 0.0 ∓ 0.0 |
| R309-W217 | 2.7 ∓ 0.4 | 2.4 ∓ 0.5 |
| R309-Y276 | 2.4 ∓ 0.6 | 2.5 ∓ 0.6 |
| R309-N221 | 1.5 ∓ 0.7 | 0.1 ∓ 0.3 |
| R312-N198 | 0.0 ∓ 0.0 | 0.0 ∓ 0.2 |
| R312-D225 | 0.0 ∓ 0.0 | 0.0 ∓ 0.0 |

Table S4: Water analysis on subunit D

[8] E. L. Wu, X. Cheng, H. Rui, K. C. Song, E. M. Dávila-Contreras, Y. Qi, J. Lee, V. Monje-Galvan, R. M. Venable, J. B. Klauda, and W. Im.

Charmm-gui membrane builder toward realistic biological membrane simulations. *Journal of computational chemistry*, 285(47):1997–2004, 2014.

[9] H. J. C. Berendsen, J. P. M. Postma, W. F. van Gunsteren, A. DiNola, and J. R. Haak. Molecular dynamics with coupling to an external bath. *The Journal of Chemical Physics*, 81(8)(3684):3684–3690, 1984.

[10] M. A. Kasimova, D. Tewari, J. Cowgill, W. Carrasquel Ursulaez, J. Lin, L. Delemotte, and B. Chanda. Helix breaking transition in the s4 of hcn channel is critical for hyperpolarization-dependent gating. *elife*, 8, 2019.
